# Changes in electrophysiological aperiodic activity during cognitive control in Parkinson’s disease

**DOI:** 10.1101/2023.10.09.561166

**Authors:** Noémie Monchy, Julien Modolo, Jean-François Houvenaghel, Bradley Voytek, Joan Duprez

**Author notes:** Correspondence to: Julien Modolo -, Full address : Laboratoire Traitement du Signal et de l’Image (LTSI), Université de Rennes 1, Campus de Beaulieu. Bât 22, 35042 Cedex - Rennes - FRANCE. These authors contributed equally to this work (Noémie Monchy, Julien Modolo, Bradley Voytek, Joan Duprez).

## Abstract

Cognitive symptoms in Parkinson’s disease (PD) are common and can significantly affect patients’ quality of life. Therefore, there is an urgent clinical need to identify a signature derived from behavioral and/or neuroimaging indicators that could predict which patients are at increased risk for early and rapid cognitive decline. Recently, converging evidence identified electroencephalogram (EEG) aperiodic activity as meaningful physiological information associated with age, development, cognitive and perceptual states or pathologies. In this study, we aimed to investigate aperiodic activity in PD during cognitive control and characterize its possible association with behavior.

Here, we recorded high-density EEG (HD-EEG) in 30 healthy controls and 30 PD patients during a Simon task. We analyzed task-related behavioral data in the context of the activation-suppression model and extracted aperiodic parameters (offset, exponent) at both scalp and source levels.

Our results showed behavioral alterations of cognitive control as well as higher offsets in patients in the parieto-occipital areas, suggesting increased excitability in PD. A small congruence effect on aperiodic parameters in pre- and post-central brain areas was also found, possibly associated with task execution. Significant differences in aperiodic parameters between the resting state, pre- and post-stimulus phases all across the scalp and cortex confirmed that the observed changes in aperiodic activity are linked to task execution. No correlation was found between aperiodic activity and behavior or clinical features.

Our findings provide evidence that EEG aperiodic activity in PD is characterized by greater offsets, and that aperiodic parameters differ depending on arousal state. However, our results do not support the hypothesis that the behavior-related differences observed in PD are related to aperiodic changes. Overall, this study highlights the importance of considering aperiodic activity contributions in brain disorders and further investigating the relationship between aperiodic activity and behavior.

## Introduction

Parkinson’s disease (PD) is a multisystem disorder characterized by motor (bradykinesia, resting tremor, postural instability and rigidity) and non-motor symptoms such as cognitive deficits.^1^ Cognitive symptoms can have a major impact on patients’ quality of life^2^ and may precede the onset of motor features.^3^ These alterations are quite heterogeneous among patients in their progression and severity.^4,5^ Currently, available treatments are ineffective in controlling cognitive symptoms and can even worsen them sometimes.^6–8^ Unfortunately, diagnostic tools to predict cognitive deterioration, or the influence of therapies on cognition, are still lacking. Therefore, there is a strong clinical need in identifying a signature derived from a number of behavioral and/or neuroimaging indicators, which could ultimately predict which patients are at increased risk of early and rapid cognitive decline.

One of the most common cognitive impairments in PD is the alteration of efficient and rapid adaptation to environmental changes. Specifically, PD patients exhibit disorders in cognitive action control (CAC), a subprocess of cognitive control that suppresses automatic responses in favor of voluntary goal-directed actions.^9^ CAC is classically studied using conflict tasks, such as the Simon task,^10^ in which participants have to respond according to the stimulus color while ignoring its location. Automatic processing of the stimulus’ spatial location can induce conflict (incongruent trials) which increases reaction times (RTs) and decreases accuracy, reflecting the so-called congruence effect.^11–13^ Most studies focusing on the effect of PD on CAC performances demonstrated a higher congruence effect on RT or accuracy in PD patients,^14–17^ although others did not replicate this result.^18,19^ To understand this divergence, some authors have used the activation-suppression model, which investigated the dynamics of CAC using distributional analyses of the congruence effect. These studies showed that PD patients tend to react automatically (greater impulsive action selection) and have difficulty suppressing the automatic activation, even when they have time to do so (impaired selective response suppression).^20–24^

The study of cognitive processes such as CAC is usually carried out using EEG with an excellent temporal resolution particularly adapted to dynamic processes. It is also arguably a more direct measure of neural activity than other neuroimaging modalities, especially regarding the measurement of neural oscillations which have been linked to behavior and notably with CAC.^25^ For instance, Singh *et al*.^26^ investigated mid-frontal theta (4-8 Hz) in PD patients during a Simon task. Results suggested that patients had an overall attenuated mid-frontal theta activity, but which was not specifically associated with changes in response conflict. This neural measure was correlated with cognitive dysfunction, supporting that cognitive failures in PD are related to attenuated cortical cognitive-control mechanisms.^27^

One difficulty in analyzing electrophysiological signals is that they typically exhibit both *periodic* and *aperiodic* properties, and these signal features are conflated.^28,29^ The periodic component, commonly referred to as neural oscillations, has received considerable attention following historical traditions, whereas the aperiodic part has only recently gained more interest and was commonly discarded or not considered. While oscillatory power is concentrated at specific frequencies, which is visible as peaks on the power spectrum,^30^ neural power spectra also exhibit a broadband 1/fχ scaling, where power decreases exponentially as a function of frequency.^31,32^ This “1/f-like” characteristic can be described by two parameters: *aperiodic offset* corresponding to the broadband offset of the spectrum, and *aperiodic exponent* defined as the χ in “1/fχ” describing the overall decreasing slope of power across frequencies.^28^ Importantly, the generative mechanisms of aperiodic activity have not been firmly established to date. Among the possible mechanisms, animal and computational models have suggested that the aperiodic exponent could be a reflection of the “excitation-inhibition” balance (the so-called “E:I ratio”) in cortical circuits.^33,34^ Recently, converging evidence points at the conclusion that changes in the aperiodic component are associated with cognitive^28,35^ and perceptual states,^36^ development,^37^ aging,^38^ and pathological conditions.^39–43^

Thus, ignoring the aperiodic component can lead to a misrepresentation and misinterpretation of physiological mechanisms.^28,37^ Furthermore, the use of *a priori* frequency bands for oscillatory analyses can result in misinterpreting aperiodic activity as periodic activity, and thus bias the power of actual physiological oscillations, or even lead to extracting power from oscillations that simply do not exist in some cases.^44^ Therefore, there is an increasing consensus on the imperative careful parametrization of spectral features to minimize its associated bias.^28^

To date, only a few studies have focused on aperiodic parameters in PD patients. Four studies have investigated local field potential signals in the subthalamic nucleus and evidenced a lower exponent during movements *versus* rest, which correlated with motor symptoms severity,^45,46^ and confirmed that aperiodic exponent can predict the motor response to subthalamic nucleus deep brain stimulation (STN-DBS).^46^ Regarding STN-DBS, a systematic increase was also found in both exponent and offset of the aperiodic spectrum 18-month follow-up after surgery in PD patients.^47^ More recently, Wiest^34^ evidenced that the aperiodic exponent of subthalamic field potentials reflects E:I ratio in PD. Five other studies were based on non-invasive scalp EEG recordings. Recent studies evidenced that PD is associated with significant increase in aperiodic activity during resting state^48,49,50^ while a decreased aperiodic activity in patients has been observed by Rosenblum *et al.*^51^ More specifically, Wang and colleagues^52^ found that aperiodic parameters were increased in PD patients after dopaminergic medication, especially in bilateral central brain regions, while Mc Keown *et al*.^50^, found that the increase in aperiodic activity in PD was independent of medication status.

To the best of our knowledge, it has still not been investigated to what extent aperiodic components are associated with behavioral alterations in PD. In this study, we focused on aperiodic activity in PD during cognitive control and tested the hypothesis that aperiodic EEG activity is related to the CAC alterations observed in PD. According to the literature, we expected to identify (i) behavioral alterations (higher congruence effect, increased impulsive action selection, decreased selective response suppression), as well as (ii) different aperiodic parameters between PD and healthy control (HC), since PD is associated with changes in cortical E:I ratio.^53^ We recorded high-density EEG (HD-EEG) from 30 HC and 30 PD patients during a Simon task. Then, we analyzed task-related behavioral data using distributional analyses and HD-EEG data by extracting aperiodic parameters at both scalp and source levels. We found behavioral alterations of CAC as well as higher offsets in PD patients in parieto-occipital brain areas, and a congruence effect on the exponent distribution at the cortex level. Aperiodic parameters were related to the overall task execution, as shown by significant changes between resting-state, pre- and post-stimulus periods. However, no correlation between behavior and EEG aperiodic parameters was found.

## Materials And Methods

### Participants

Thirty HC (15 males, 15 females) aged between 45-70 years (mean = 61.7, sd = 7.3) and 30 patients (15 males, 15 females) diagnosed with idiopathic PD,^54^ aged between 45-73 years (mean = 60.4, sd = 7.3) were enrolled in this study. One patient was excluded due to a technical acquisition issue during an experimental session. Thus, all subsequent analyses were performed on 30 HC and 29 PD patients (Table 1). HC and patients did not significantly differ in age, sex or education.

**Table 1.** Demographic characteristics of participants. Data are presented as mean (standard deviation)

|  | HC (n = 30) | PD (n = 29) | t-test |
| --- | --- | --- | --- |
| Age (years) | 61.7 (7.3) | 60.4 (7.3) | t = 0.77; p = 0.44 |
| Sex | 15 : 15 | 14 : 15 | - |
| Education (years) | 13.5 (3.6) | 12.5 (3.2) | t = 1.19; p = 0.24 |
| MoCA (/30) | 28 (1.9) | 26.6 (2.0) | t = 2.72; p <0.01 |
| Disease duration | - | 12.1 (4.2) | - |
| LEDD (mg/day) | - | 1348.0 (462.0) | - |
| UPDRS III - ON | - | 10.0 (5.7) | - |
| UPDRS III - OFF | - | 36.8 (12.5) | - |
| Hoehn & Yarh - ON | - | 1.1 (0.8) | - |
| Hoehn & Yarh - OFF | - | 2.0 (0.4) | - |
| Schwabb & England - ON | - | 93.0 (5.6) | - |
| Schwabb & England - OFF | - | 76.5 (10.3) | - |

MoCA: Montreal Cognitive Assessment Scale; LEDD: Levodopa Equivalent Daily Dose; UPDRS: Unified Parkinson's Disease Rating Scale

All patients were recruited from the Neurology Department of Rennes University Hospital (France). They were on dopaminergic medication, measured by L-dopa equivalent daily dose (LEDD), for all parts of the experiment (Table 1). Patients with DBS devices were not included. All HC were recruited from the general population by advertising. Participants had normal or corrected-to-normal vision.

All participants underwent a neuropsychological assessment that revealed no major cognitive impairment (all Montreal Cognitive Assessment Scale, MoCA scores > 22)^55^ nor severe neurocognitive disorder according to the Diagnostic and Statistical Manual of Mental Disorders - V (DSM-V). All participants were also free from moderate or severe psychiatric symptoms and present or past neurological pathology (other than PD for patients). The severity of patients’ motor symptoms was assessed using the Unified Parkinson’s Disease Rating Scale (UPDRS) III scale^56^ (Table 1).

This study was conducted in accordance with the declaration of Helsinki and was approved by a national ethics committee (CPP ID-RCB: 2019-A00608-49; approval number: 19.03.08.63626). After a complete description of the study, all participants gave their informed written consent.

### Task design and procedure

Participants were asked to perform a color version of the Simon task^10^ to assess CAC. They were asked to press a right or left button, as quickly and accurately as possible, according to the color of the circle while ignoring its right or left location on the computer screen (color/side mapping was counterbalanced across participants). Participants had to respond within 1000 ms after stimulus offset. In congruent trials, the side of presentation of the circle matched with the side of the button press associated with the color, facilitating the response of the subject. In incongruent trials, color and location didn’t match and activated conflicting responses (Supplementary Fig.1). Participants first performed a training session composed of 60 trials and then an experimental phase organized in 10 blocks of 60 trials. Each block contained 30 congruent and 30 incongruent randomized trials. In total, 600 trials were performed with 300 congruent and 300 incongruent trials. For further details, see.^57^

## HD-EEG recording and processing

### Recording

EEG data was recorded using a HD-EEG system (EGI, Electrical Geodesic Inc., 256 channels) with a sampling frequency of 1000 Hz, an electrode impedance maintained below 25 kΩ, the Cz electrode used as the reference and a ground close to Pz. Electrodes on the net were placed using the standard 10-10 geodesic montage. Most jaw and neck electrodes were removed due to excessive muscular artifacts, resulting in a total of 199 exploitable electrodes. The EEG recording consisted in a 5-minutes resting state period with eyes closed followed by the training.

### Preprocessing

All EEG preprocessing and subsequent analyses were performed manually using the Brainstorm toolbox^58^ in Matlab (MathWorks®, USA). First, DC offset removal was applied. Second, a notch filter (50 Hz) and a band-pass filter (1-90 Hz) were applied (Finite Impulse Response; FIR filter using Kaiser window). Third, signals were visually inspected and bad channels were removed before being interpolated using Brainstorm’s default parameters. Fourth, Independent Component Analysis (ICA, *jade* method) was used to remove eye blinks and muscle artifacts following visual inspection of the independent components. Fifth, signals were segmented into 4-second epochs for the resting-state period and from −700 ms to 1200 ms relative to the stimulus onset during the task. Finally, a visual inspection was performed to manually reject remaining bad epochs. As a result, an average of 59 (sd = 14) resting-state epochs and 326 (sd = 13) correct trials were studied per subject. Only correct congruent and incongruent trials were studied, since they are associated with efficient control.

After preprocessing, several steps were applied to extract aperiodic parameters at both scalp and source levels, as summarized in Fig. 1 and explained below.

**Figure 1.**
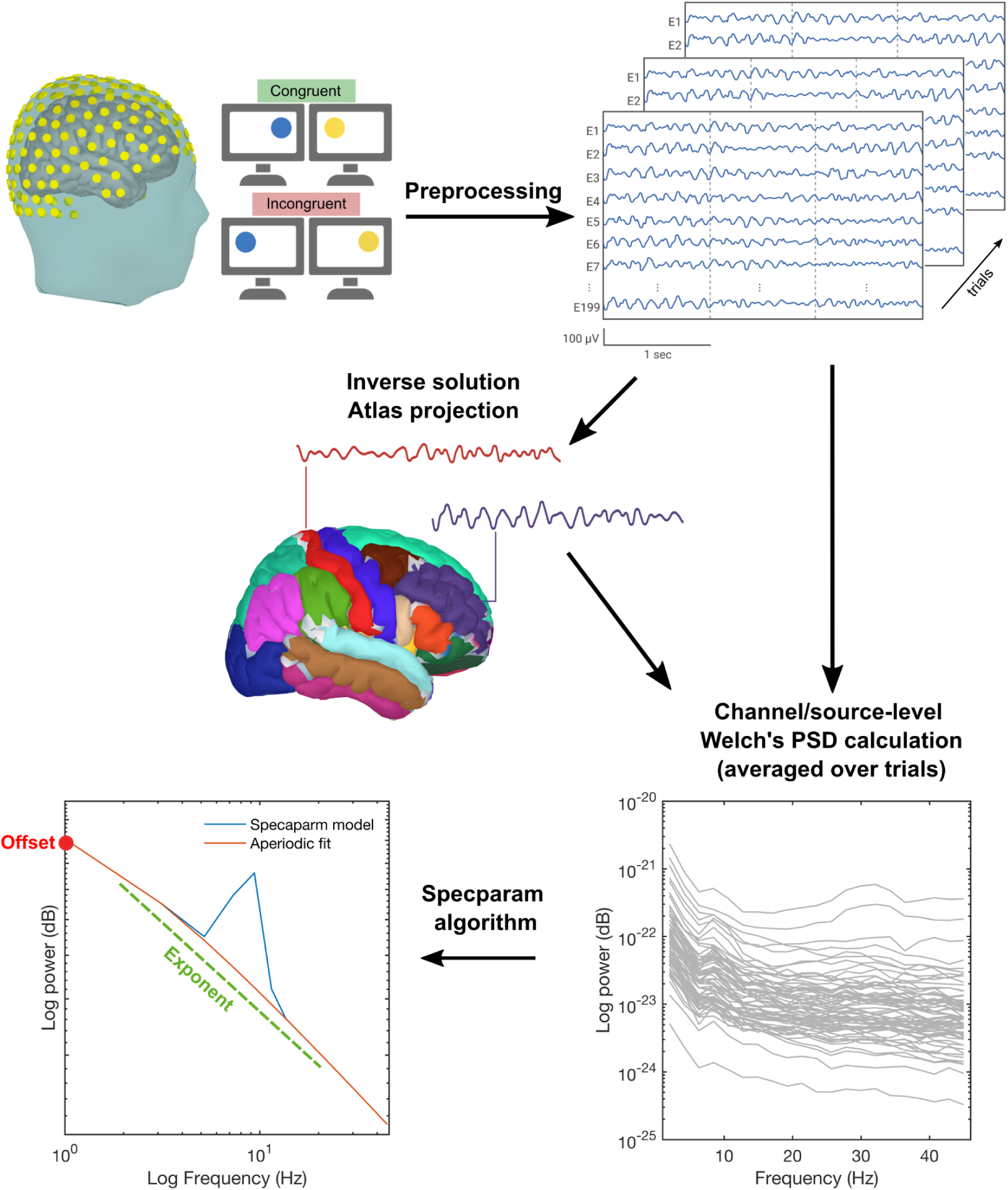
Overview of the pipeline analysis of HD-EEG data. First, HD-EEG data was recorded during the Simon task (only correct trials were considered for the post-stimulus part). After preprocessing, PSD was estimated on the resting-state period, on the prest-stimulus part (−500 to 0 ms) and on the post-stimulus part (0-1000 ms) for each epoch. The spectra obtained were averaged by subject and condition for the post-stimulus part and only by subject for resting state and prestimulus period. Finally, we applied the *specparam* algorithm to extract the aperiodic parameters for each averaged spectrum.

### Source reconstruction

A realistic head model was used along with the position of the electrodes. In this study, we used the Boundary Element Method (BEM) head model fitted to the International Consortium for Brain Mapping Magnetic Resonance Imaging (ICBM MRI) template,^59^ and the OpenMEEG toolbox.^60^ The EEG inverse problem was solved with the weighted minimum norm estimate (wMNE) method.^61^ We used the Desikan-Killiany Atlas parcellation (68 regions of interest; ROIs)^62^ to project the cortical sources, and constrained their orientation normally to the cortical surface.^63^ We used a noise covariance matrix computed from our 700 ms pre-stimulus baseline and set the signal to noise ratio (SNR) and the depth weighting to default values.

## Parameterization of the spectral data

Power spectrum density (PSD) was obtained for each epoch using Welch’s method (window length of 0.5 second and window overlap ratio of 50%), which is a necessary prerequisite for the subsequent estimation of aperiodic parameters. PSD was calculated on the resting-state period, on the pre-stimulus part, i.e., from −500 to 0 ms before the stimulus appeared on the screen, and on the post-stimulus part, i.e., from 0 to 1000 ms after the stimulus appeared on the screen. The obtained spectra were averaged by subject and condition for the post-stimulus part, and by subject only for the resting state and pre-stimulus part. For each resulting power spectrum, we applied the spectral parameterization algorithm across the 1–40 Hz frequency band (*specparam* toolbox^64^ version 1.0), which considers the PSD as a linear combination of two different types of components: aperiodic and periodic (oscillatory) components.^28^ The aperiodic component consisted of the exponent (overall spectrum slope) and offset (intercept of the curve) parameters. In order to optimize parameters of the *specparam* algorithm to our data, we compared the goodness-of-fit metrics of several *specparam* models and selected the model with the best fitting metrics: (channel-averaged R^2^ = 0.94; channel-averaged mean squared error, MSE = 4.5^-3^). The optimal parameters were: peak width limits: [0.5-6]; max number of peaks: 4; minimum peak height: 1.0; peak threshold: 2.0 and aperiodic mode = ‘fixed’. Aperiodic parameters were estimated at both the scalp and the cortical ROI level.

## Statistical analysis

All statistical analyses were conducted in R v.4.1.3.^65^ implemented with the *tidyverse* and *lme4* package.^66,67^ For all analyses, we chose the standard significance threshold of p = 0.05.

### Behavioral responses

For each participant and for each trial, RT and accuracy scores were extracted. To estimate the congruence effect, we compared congruent and incongruent RT of correct responses as well as accuracy. Then, we also compared these data between groups to estimate the impact of PD. In light of the activation-suppression model,^68^ we further analyzed behavioral data with distributional analyses.^69,24^ First, impulsive action selection (incongruent accuracy of the fastest trials) was investigated using conditional accuracy functions (CAFs) displaying accuracy rate against the RT distribution for each condition and group. For each participant, RTs were rank-ordered and split into seven bins containing an equal number of trials, and mean accuracy was then plotted for each bin. The activation-suppression model postulates that incongruent accuracy of the first bin (fastest trials) informs about impulsive action selection. Second, we used delta plots to assess the dynamics of selective response suppression (slope value of the congruence effect for the slowest trials). Delta plots display the mean congruence effect (incongruent RT - congruent RT) as a function of the RT distribution of correct responses, splitted into seven bins, as we did for CAFs. The activation-suppression model postulates that the slope between the two last bins of delta plots informs about the selective response suppression.

The effect of congruence and group on RT and accuracy were analyzed using linear and non-linear mixed models, respectively. These models are composed of two fixed factors: group and congruence, and a random effect of subject and condition by subject was added. Overall, 551 (sd = 81) correct trials on average per subject were kept for behavioral analyses due to the removal of errors, absence of response within 1000 ms or remaining artifacts. RT were log-transformed for increased compliance with the model’s assumptions (homogeneity of variance and normal distribution of residuals model).

Fixed effects significance were computed through the Anova function of the {*car*} package^70^ that calculates type II Wald chi-square tests. Marginal (mR^2^) and conditional (cR^2^) were calculated using the {MuMin} package.^71^ These models offered the possibility to exploit the whole dataset, avoiding the loss of statistical power due to averaging data, and also enabled taking into account interindividual variability and unbalanced data.^72^

Group effect on the average first bin of CAF and on the last slope values of delta plots were estimated with Welch’s t-test, since only one averaged value exists for each subject, and are reported with effect size (Cohen’s d).

### EEG parameters

Following the extraction of aperiodic parameters, we first performed analyses on the post-stimulus part (0 to 1000 ms after the stimulus onset) on both scalp and cortex levels. Within each level, we studied the aperiodic parameters averaged over all the electrodes/ROIs and for each electrode/ROI. Group and congruence effects on aperiodic offset and exponent were assessed using two-way repeated measures ANOVAs where the between-subject factor variable was the group and the within-subject factor variable was the congruence. Classical ANOVAs were chosen over mixed models here because mixed-modeling only converges when several repetitions by factor combination are available, which was not the case of the aperiodic parameters as compared to behavioral data. Assumptions were checked for each test (outliers and normality assumption). Paired student t-tests were performed as post-hoc tests where the effect was significant. All post-hoc tests were FDR (False Discovery Rate)-corrected.

Then, we analyzed group and period effects (resting state vs. pre-stimulus period vs. post-stimulus period) on aperiodic parameters in order to verify if aperiodic parameters were different when participants were performing the task compared to resting state and pre-stimulus periods. We applied the same pipeline analysis as described above except that the within-subject factor variable of the two-way repeated measures ANOVAs was the period (resting-state, pre-stimulus or post-stimulus part).

### Correlations between behavior and EEG parameters

Last, we tested whether behavioral variations were associated with aperiodic parameters. To achieve this, we used Spearman correlations between behavioral responses as dependent variables (RT, accuracy, first bin of CAF, last slope of delta plots) and EEG parameters from *specparam* as independent variables (offset, exponent). We then applied a FDR correction to account for multiple-comparisons tests. We also tested correlations between aperiodic parameters and clinical characteristics of the PD population (age, disease duration, UPDRS-III ON/OFF scores, LEDD).

### Code availability

The channel file, and all the Matlab and R codes used for preprocessing and data analysis are publicly available at (https://github.com/noemiemonchy/PD-SIMON).

## Results

### Behavioral analysis

The congruence effect classically reported in the literature was observed, as indicated by longer RTs (*χ*^2^ = 451.9, p < 0.0001; mR^2^ = 0.06; cR^2^ = 0.42) and more errors (*χ*^2^ = 163.1, p < 0.0001; mR^2^ = 0.18; cR^2^ = 0.3) in the incongruent versus congruent situation (Fig. 2A-2B). Thus, participants were slower and made more errors overall during conflict. PD patients were less accurate than HC regardless of congruence (*χ*^2^ = 18.1, p < 0.0001), but had a similar RT (*χ*^2^ = 1.3, p = 0.25). The congruence effect was not significantly different between groups, both for RT (*χ*^2^ = 0.25, p = 0.62) and accuracy (*χ*^2^ = 0.23, p = 0.63). This suggests that conflict resolution, based only on the analyses of overall congruence effect, was not different between the two groups.

**Figure 2.**
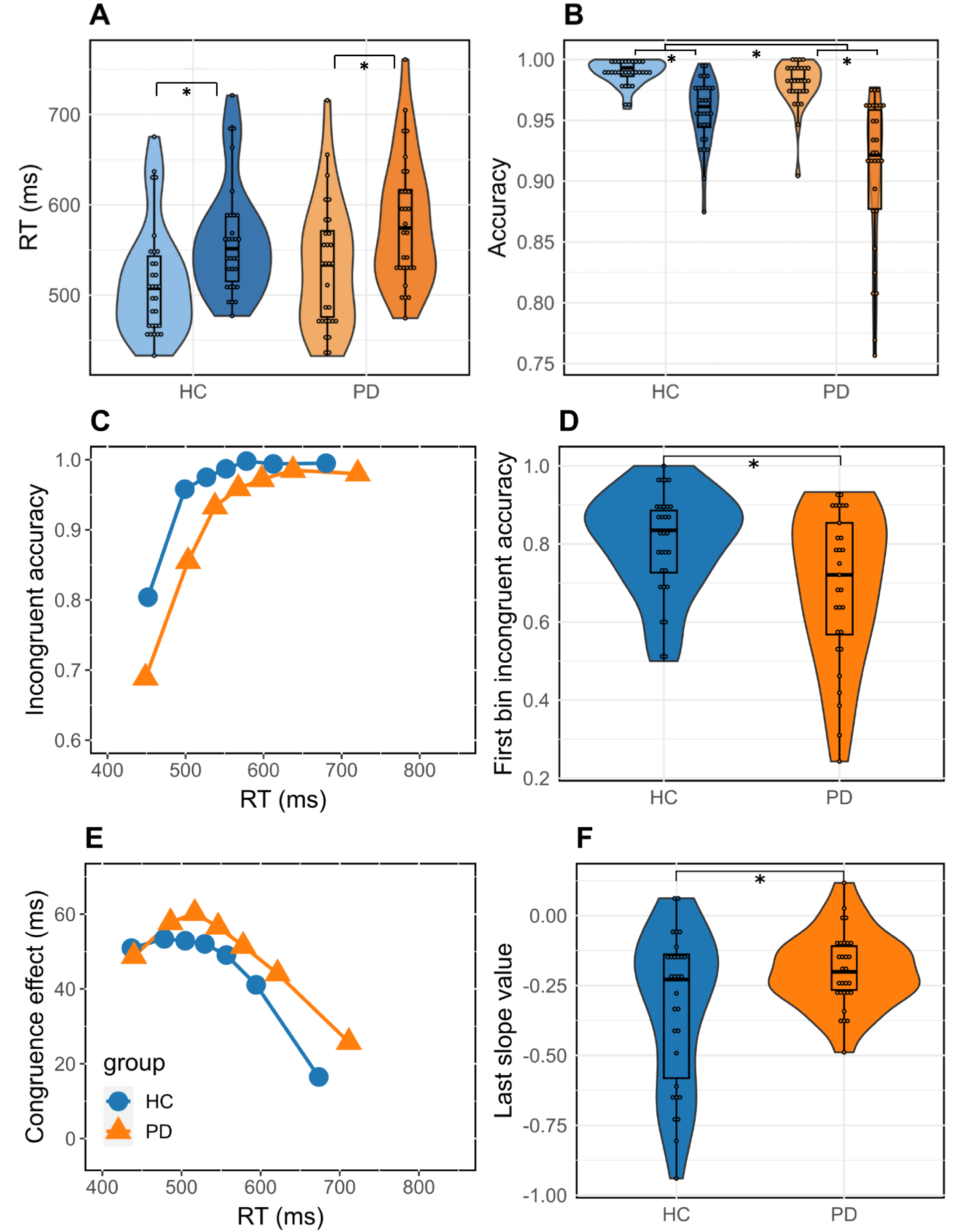
Subject-averaged reaction time (A) and accuracy (B) as a function of congruence in both groups. The lightest color corresponds to congruent trial values, while the darkest one corresponds to incongruent trial values. CAFs for the incongruent situation are plotted against RT distribution as a function of group. Impulsive action selection is denoted by violin plots showing accuracy of the first incongruent bin (D) by group. Delta plots showing changes in the congruence effect as a function of RT distribution in both groups (E). The strength of selective inhibition is represented by violin plots showing the value of the last slope of the delta plots in both groups (F). *p < 0.05

CAFs revealed the typical pattern of lower accuracy for the fastest RTs (Fig. 2C). Accuracy of the first bin (the fastest responses) in the incongruent condition (Fig. 2D) was significantly lower in PD patients in comparison with HC (t = 2.67, p = 0.01; Cohen’s d = 0.70), reflecting a higher impulsive action selection in the patients group.

Delta plots also displayed the typical decreasing pattern of the congruence effect with RT associated with the Simon task (Fig. 2E). PD patients had a significantly flatter slope, indicating reduced ability in inhibiting automatic responses (Fig. 2F; t = −2.44, p = 0.02; Cohen’s d = −0.63).

### EEG analysis

#### Group and congruence effects on aperiodic parameters

##### Scalp level

We first took a global approach, averaging the exponent and offset values over all the electrodes for each subject (Fig. 3A-B). The electrode-averaged exponent and offset did not differ according to congruence (exponent : F(1,57) = 0.23; p = 0.63; offset : F(1,57) = 0.044; p = 0.834). However, offset values significantly differed between groups : PD had higher aperiodic offsets (F(1,57) = 6.36; p = 0.014) than HC, while electrode-averaged exponent values did not differ between groups (F(1,57) = 0.091; p = 0.76).

**Figure 3.**
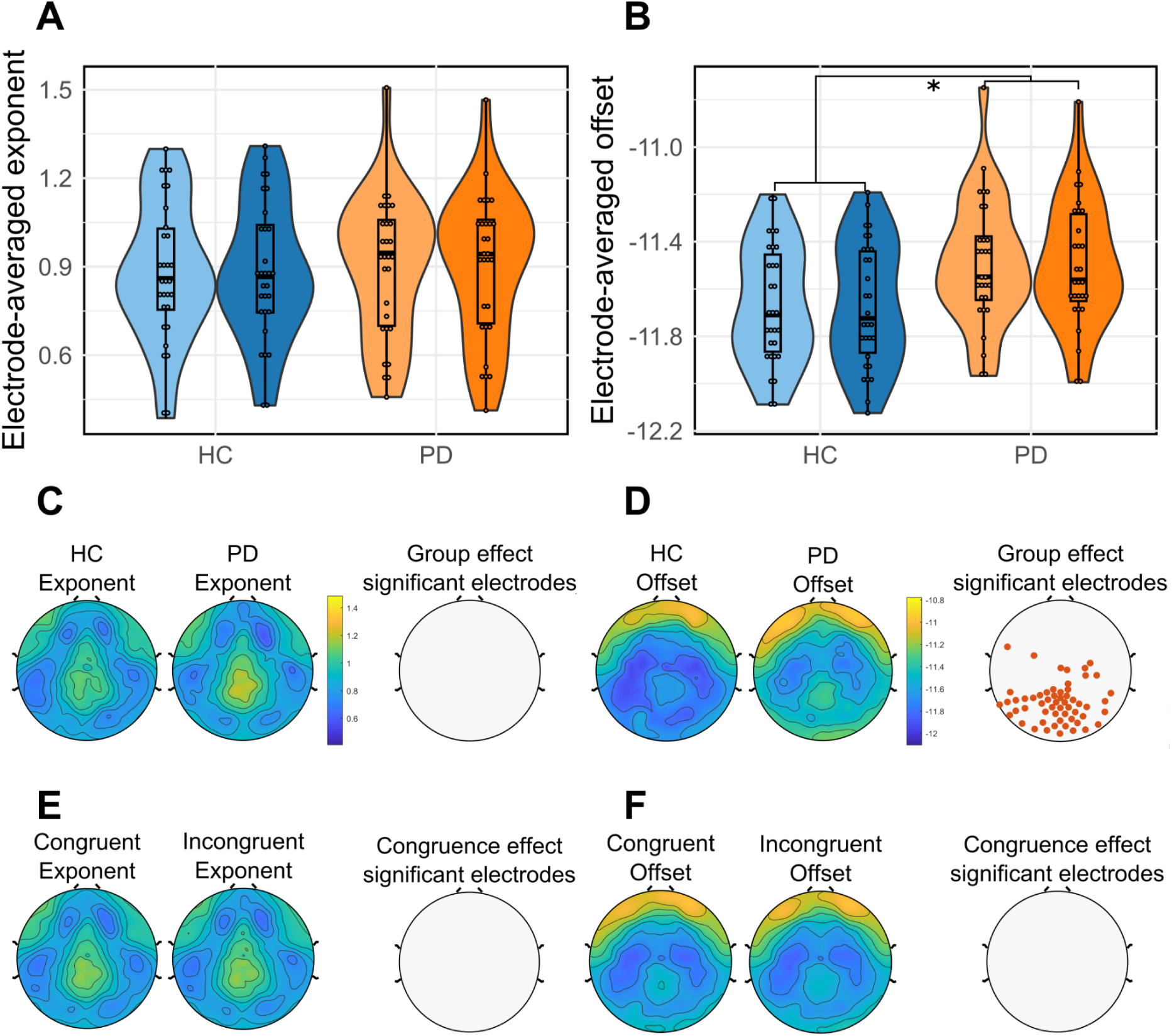
Violin plots showing the distribution of electrode-averaged exponent (A) and offset (B) values by condition and for each group. The lightest color corresponds to congruent trial values, while the darkest one corresponds to incongruent trial values. Spatial topographies of the aperiodic exponent (C) and offset (D) values across the scalp regardless congruence for HC and PD patients. Comparisons between groups were performed with ANOVAs for each electrode (FDR-corrected). Electrodes with significant differences in aperiodic parameters between HC and PD patients are represented with red dots. Spatial topographies of the aperiodic exponent (E) and the offset (F) values across the scalp regardless of the group.

Then, we analyzed the distribution of aperiodic parameters for all electrodes across the scalp for each group regardless of condition (Fig. 3C-D). We observed that the distribution pattern of the offset and exponent values seemed similar between HC and PD patients. Regarding exponent distribution, no significant differences in values have been shown between groups (Fig. 3C). Aperiodic offsets in PD were significantly higher as compared to HC (Fig. 3D) over a large majority of parieto-occipital electrodes (Fig. 3D).

Distribution of aperiodic parameters across the scalp was also estimated for each condition regardless of group (Fig. 3E-F). We found no significant differences in aperiodic offset and exponent between conditions.

##### Source level

The ROI-averaged exponent and offset did not differ according to congruence (exponent : F(1,57) = 0.065; p = 0.8; offset : F(1,57) = 0.101; p = 0.752). However, ROI-averaged offset values significantly differed between groups: PD had higher aperiodic offsets (F(1,57) = 6,17; p = 0.016) than HC, while ROI-averaged exponent values did not differ between groups (F(1,57) = 0.705; p = 0.405).

Distribution of aperiodic parameters for all ROIs was estimated for each group regardless of condition (Fig. 4C-D). We observed that the distribution pattern of the offset and exponent values seemed similar between HC and PD patients. Regarding exponent distribution, no significant differences in values have been evidenced between groups (Fig. 4C). Aperiodic offsets in PD were overall higher as compared to HC over left parieto-occipital regions and on both temporal lobes (Fig. 4D; Supplementary Table 1).

**Figure 4.**
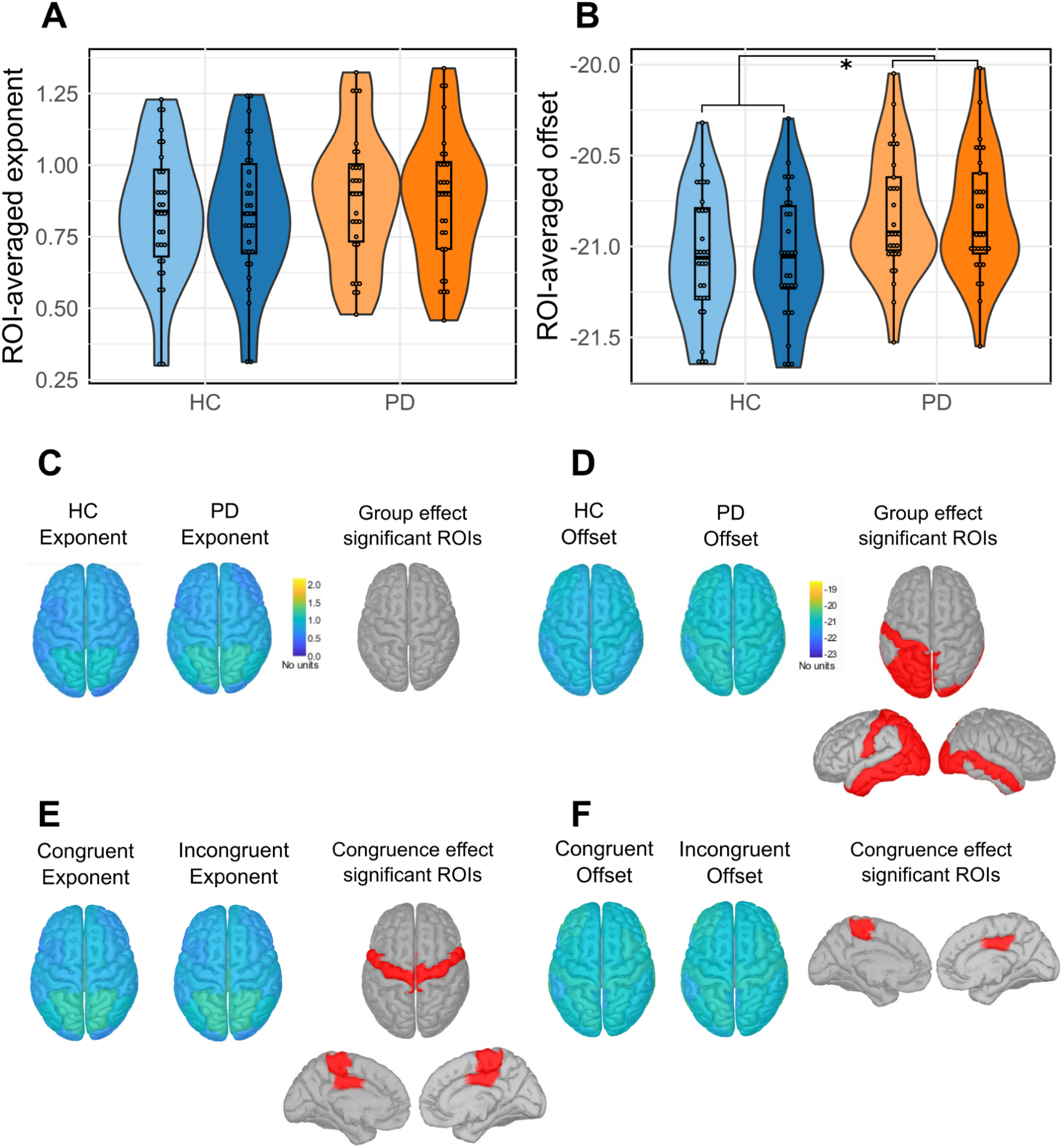
Violin plots showing the distribution of ROI-averaged exponent (A) and offset (B) values by condition and for each group. The lightest color corresponds to congruent trial values, while the darkest one corresponds to incongruent trial values. Spatial topographies of the aperiodic exponent (C) and offset (D) values across the cortex regardless of congruence for HC and PD patients. Comparisons between groups were performed with ANOVAs for each ROI (FDR-corrected tests). ROIs with significant differences in aperiodic parameters between group or condition are represented in red. Spatial topographies of the aperiodic exponent (E) and offset (F) values across the cortex regardless of the group.

Distribution of aperiodic parameters across the cortex was also estimated for each condition regardless of group (Fig. 4E-F). The distribution pattern of aperiodic parameters seemed overall similar between conditions too. However, significant differences in exponent parameters were found in precentral areas (Supplementary Table 1). Offset significantly differed in the right posterior cingulate and left precentral medial area (Supplementary Table 1). Aperiodic exponents in incongruent conditions were greater than in congruent trials but the effect size associated with these differences were very low (congruent: mean = 0.85, sd = 0.4; incongruent: mean = 0.86, sd = 0.4; size effect: 6.7^-3^). Same is found in offset values (congruent: mean = −21.03, sd = 0.6; incongruent: mean = −20.9, sd = 0.6; size effect: 0.149).

### Task period effect on aperiodic parameters

We applied the same approach to test whether aperiodic parameters were different between resting-state and during the task (pre- and post-stimulus) to check that our results didn’t only reflect baseline aperiodic activity.

#### Scalp level

Electrode-averaged exponents did not significantly differ between groups (F(1,57) = 1.48; p = 0.229), but significantly differed according to the task period (F(2,114) = 82.47; p < 0.0001; Fig. 5A). In addition to significant differences according to the task period (F(2,114) = 74.018; p < 0.0001), electrode-averaged offsets also differed between groups (F(1,57) = 14.71; p = 3.15^-4^; Fig. 5B).

**Figure 5.**
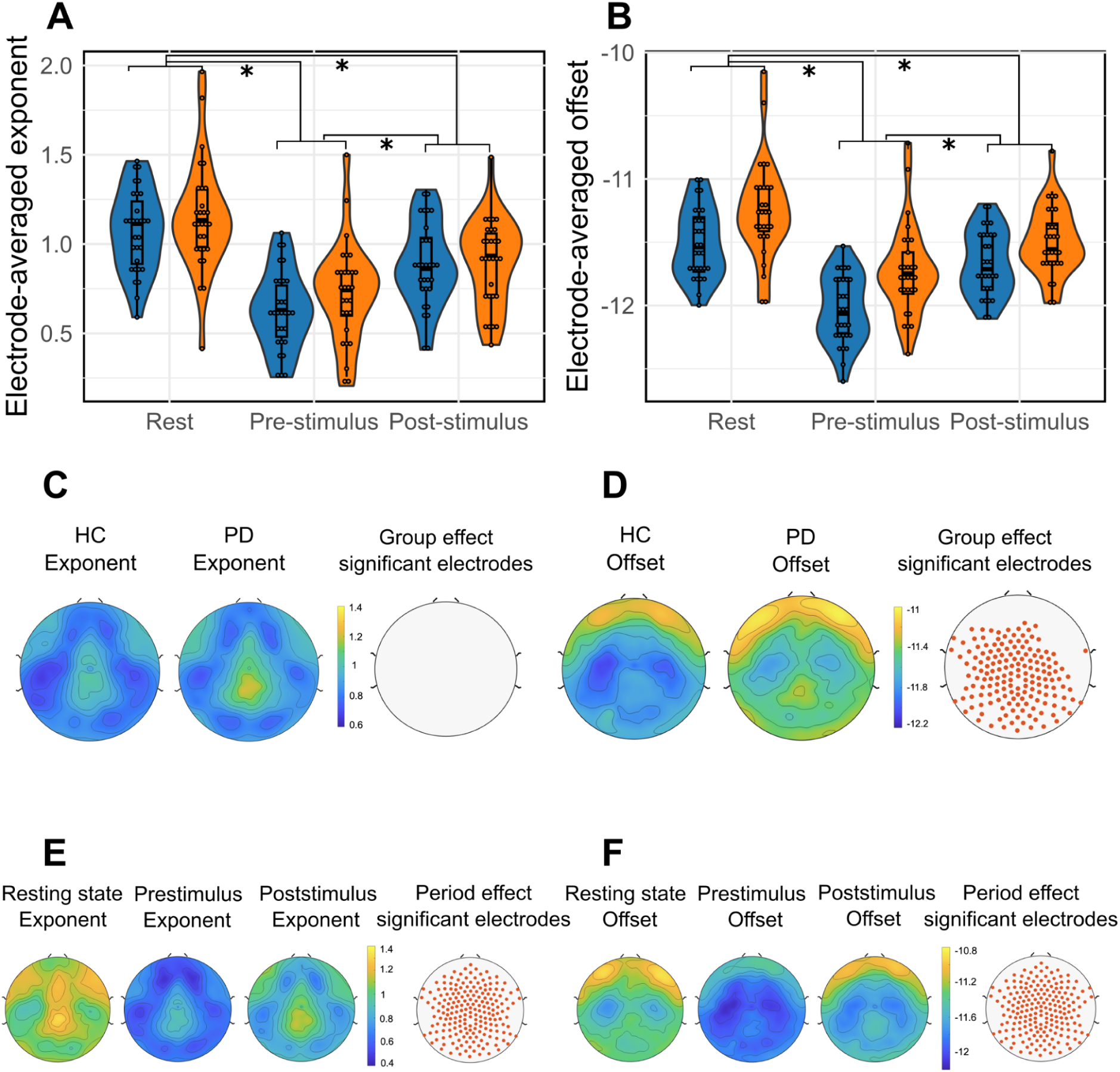
Violin plots showing the distribution of electrode-averaged exponent (A) and offset (B) values by group (blue for HC, orange for PD patients) and for each task period based on all electrodes. Spatial topographies of the aperiodic exponent (C) and offset (C) values across the scalp regardless of task period for HC and PD patients. Comparisons between groups were performed with ANOVAs for each electrode (FDR-corrected tests). Electrodes with significant differences in aperiodic parameters between HC and PD patients are represented with red dots. Spatial topographies of the aperiodic exponent (E) and the offset (F) values across the scalp regardless of the group.

The distribution of exponent values across the scalp for each group regardless of task period showed no significant group effect (Fig. 5C). However, the task period significantly differed regardless of group on all electrodes across the scalp (Fig. 5E). *Post-hoc* tests evidenced that exponents were greater (more negative slope) during the resting-state than during the post-stimulus period (p = 1.13^-8^), and even more so than during the pre-stimulus period (p = 2.09^-17^; see Supplementary Fig. 2A).

Distribution of the offset across the scalp was estimated for each group regardless of the task period (Fig. 5D). Offsets were higher in PD patients over a large majority of electrodes across the scalp, except in frontal areas, especially on the right (Fig. 5F). Regarding the offset distribution across the scalp for each task period regardless of the group, offsets were significantly different according to the period on the whole scalp. *Post-hoc* comparisons showed that offsets during the resting-state period were higher than offsets during the pre-stimulus period on all electrodes (p = 6.5^-17^) and higher than offsets during the post-stimulus period on a large majority of electrodes across the scalp (p = 1.56^-5^). Moreover, post-stimulus offsets were significantly higher than pre-stimulus offsets on all electrodes across the scalp (p = 8.78^-9^; see Supplementary Fig. 2B)

#### Source level

ROI-averaged exponents did not significantly differ between groups (F(1,57) = 1.13; p = 0.292), while the task period had a significant effect on exponents (F(1.24,71.7) = 122.46; p < 0.0001). The group had a significant effect on ROI-averaged offsets (F(1,57) = 6.079; p = 0.017). ROI-averaged offsets also significantly differed according to the task period (F(1.34,76.23) = 122.46; p < 0.0001).

The distribution of exponent values across the scalp for each group regardless of task period showed no significant group effect (Fig. 6C). However, the distribution of ROI-averaged exponents according to the task period regardless of the group showed a significant effect of the task period on all ROIs (Fig. 6E; Supplementary Table 1). Similar to the electrode-level results, *post-hoc* tests evidenced that exponents were greater (more negative slope) during the resting-state than during the post-stimulus period (p = 2.33^-5^) and even more so than during the pre-stimulus period (p = 7.05^-20^; Supplementary Fig. 3A).

**Figure 6.**
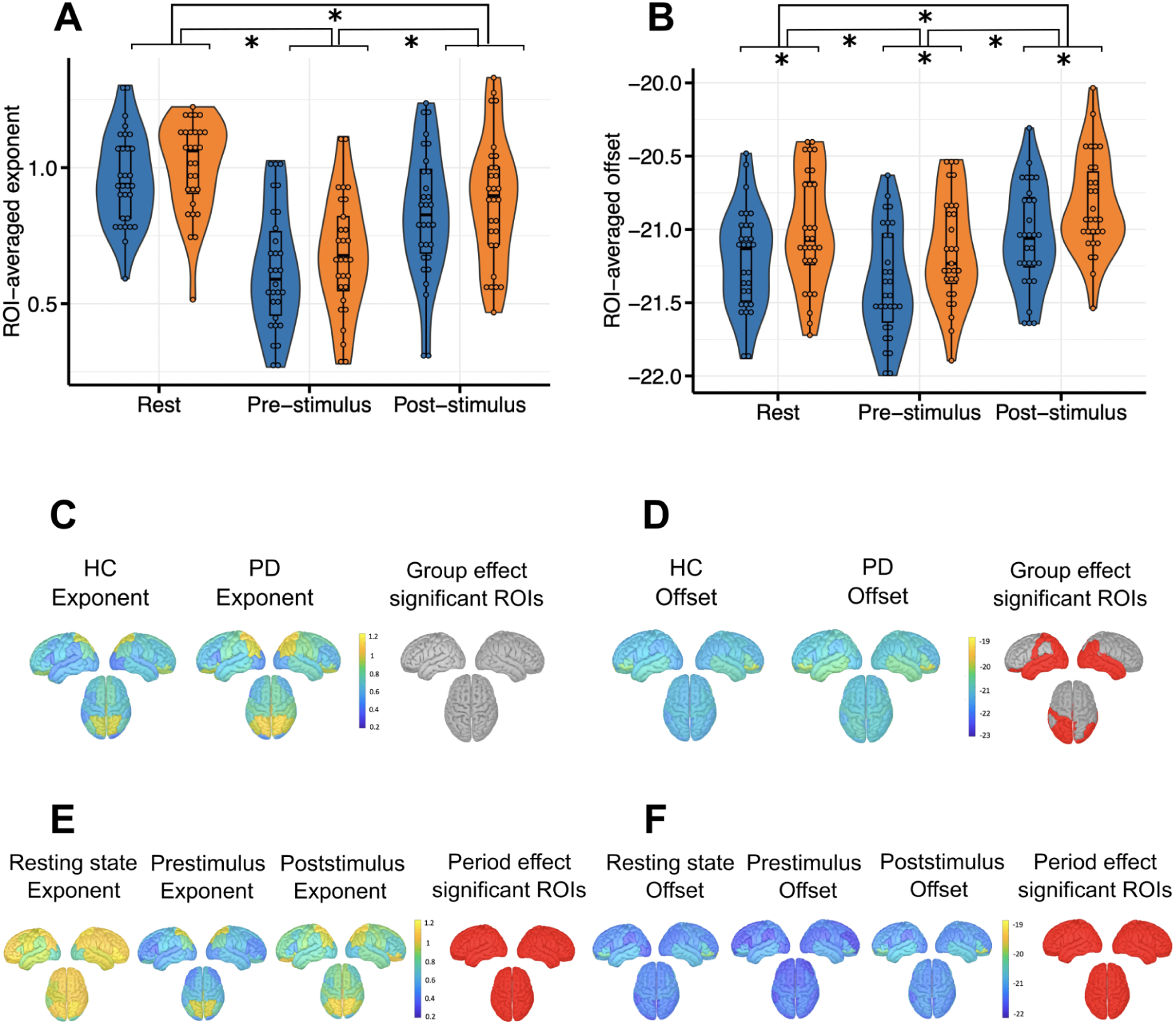
Violin plots showing the distribution of ROI-averaged exponent (A) and offset (B) values by group and for task period. Spatial topographies of the aperiodic exponent (C) and offset (D) values across the cortex regardless of task period for HC and PD patients. Comparisons between groups were performed with t-tests for each ROI (FDR-corrected tests). ROIs with significant differences in aperiodic parameters between group or condition are represented in red. Spatial topographies of the aperiodic exponent (E) and offset (F) values across the cortex regardless of the group.

Distribution of the offset across the cortex was estimated for each group regardless of the task period (Fig. 6C). Offsets were higher in PD patients over occipital and left parietal cortical regions as well as in both temporal lobes (Fig. 6D; Supplementary Table 1). Regarding the offset distribution across the cortex for each task period regardless of the group, offsets were significantly different according to the period on the whole cortex. *Post-hoc* tests showed that offsets during the post-stimulus period were greater than resting-state offsets (p = 9.36^-6^) which were themselves greater than offsets during the pre-stimulus period (p = 2.6^-5^; Supplementary Fig. 3B)

### Correlation analysis

Correlation analyses did not find any significant association between behavioral or clinical variables and aperiodic parameters.

## Discussion

The aim of this study was to test our hypothesis that aperiodic activity in the EEG power spectrum differs in PD in comparison with HC during CAC, and that these aperiodic changes are linked to the CAC alterations observed in PD. Several key findings emerged from this study. First, we identified an alteration of CAC in PD, as was expected from previous studies. Second, HD-EEG data evidenced that, in the post-stimulus period, the aperiodic offsets were higher in PD patients in parieto-occipital areas at both scalp and cortex levels. In contrast, no group effect was found on the exponent distribution, except a congruence effect in motor areas. In addition, aperiodic parameters were both significantly different according to the task period: changes in offsets and exponents were found between resting-state, pre- and post-stimulus period. Finally, no correlation between aperiodic activity and behavior or clinical characteristics of the PD population was found, thereby not supporting our hypothesis.

### Cognitive alteration in Parkinson’s disease

Our results at the Simon task are consistent with the well-known CAC alterations in PD. PD patients, as compared to HC, had an overall lower accuracy, as well as higher impulsive action selection and decreased strength of selective response suppression. No higher congruence effect or higher overall RT were observed in PD. Changes in the dynamic expression of CAC regarding impulsive action selection^22,24^ or suppression^20–23^ have been indeed documented. The lack of a greater congruence effect on overall data may be due to several factors, such as the task settings,^73^ or according to the population characteristics that vary a lot among studies. Indeed, the PD patients included in our study exhibited mild to moderate cognitive alteration.^74^ However, it has been found that the extent of cognitive disorders in PD is quite heterogeneous,^75^ and symptom severity could be associated with a higher congruence effect or speed accuracy trade off. ^21,76^

### Group effect on aperiodic parameters

Our EEG analysis indicated an approximately symmetrical spatial distribution of the aperiodic parameters in HC and PD patients, both at the scalp and source level. These topographies are similar to those already observed in the literature in control^77^ and in patients.^52^ This suggests that aperiodic activity distribution pattern is preserved in PD patients.

We observed significantly higher offsets in PD patients *versus* control in the parietal-occipital regions. This group effect was found at both averaged and specific scales at scalp and cortex levels. Our results also evidenced that aperiodic offsets were significantly higher in PD patients in comparison to HC during the post-stimulus phase but also more globally, regardless of task period. This result is in line with previous studies demonstrating greater offsets in PD patients during resting-state.^49,50,52^ More particularly, Wang *et al.*^52^ found that aperiodic offsets were greater in PD patients *versus* control in parietal-occipital midline regions during resting-state. Some studies suggested that aperiodic offset is linked to the rate of neural spiking,^78,79^ involving that these significant differences driven by aperiodic offset could be caused by increased neuronal spiking at the cortical level in PD. This is supported by previous studies showing that neural spiking rate in PD increased with motor symptom severity and disease progression.^80^

However, no significant difference has been evidenced between HC and PD regarding exponent distribution, which contrasts with what has already been observed in the literature. Indeed, previous studies on aperiodic activity showed that exponents were significantly higher in PD patients *versus* control during resting-state.^48–52^ Recently, Helson *et al*.^48^ demonstrated that the exponent was significantly larger in PD patients than in HC in all cortical areas except frontal ones, with stronger changes in sensory and motor areas. Physiological evidence has demonstrated that the aperiodic exponent can reflect the balance between excitation and inhibition.^33,81^ A flatter exponent is assumed to translate an increase in E:I ratio, while a steeper exponent is assumed to reflect a decreased E:I ratio. Moreover, previous studies also showed that impairments in dopaminergic and GABAergic neuronal activity are prominent in PD, leading to an imbalance in E:I ratio.^82–84^ Our exponent results could be explained by the clinical criteria of our PD population. Indeed, our PD patients were younger^50,51^ and had a less severe disease in comparison to the PD population studied previously.^48,50,52^ We can hypothesize that slight to moderate symptoms are less associated with an imbalance in E:I ratio, and consequently with more modest changes in aperiodic exponent.

### Congruence effect on aperiodic parameters

A congruence effect on central and posterior cingulate areas was only found at the cortical level. This result was not visible on a global scale, which underlines the very localized aspect of the congruence effect on aperiodic parameters. Aperiodic parameters were greater in incongruent condition, however the effect size was low for both aperiodic parameters, thus preventing strong conclusions. Very few studies have investigated aperiodic activity in the context of experimental cognitive tasks, and any with PD patients, which limits the comparison of our results with the literature. Only one study focused on aperiodic neural activity during metacontrol, where Zhang *et al*.^85^ examined whether aperiodic activity reflects metacontrol analyzing EEG and behavioral data of HC performing a Simon Go/NoGo task. They evidenced higher aperiodic parameters in NoGo trials compared with Go trials in incongruent trials, as compared to congruent ones. However, McSweeney and colleagues^86^ investigated changes in aperiodic activity during early adolescence in a healthy population. They used an experimental task to assess the ability to suppress inappropriate responses with a Flanker task, and found no significant effect of condition on aperiodic parameters. In our study, aperiodic parameters were greater in motor areas in incongruent conditions as compared to congruent ones. This difference can be associated with the motor execution of the task: incongruent conditions requiring greater motor control could be associated with an increase in aperiodic activity.

### Task period effect on aperiodic parameters

Our study showed significant changes in aperiodic parameters regarding the task period: both offsets and exponents were significantly different between resting-state, pre- and post-stimulus period over a large majority of electrodes and brain areas. This result is consistent with those obtained in the study focusing on the link between aperiodic activity and metacontrol. Zhang and his colleagues^85^ found that EEG power spectrum aperiodic activity reflected metacontrol states and particularly dynamic adjustments of mentacontrol states to task demands. It has also been demonstrated that the aperiodic exponent among children reflects distinct cognitive processes.^87^ Moreover, a study from Lendner *et al*.^88^ also demonstrated that the aperiodic exponent delineates wakefulness from anesthesia, rapid eye movement (REM) and non-REM sleep. In that sense, different arousal and cognitive states associated to the distinct task periods could explain these changes in both aperiodic parameters. These results suggest that aperiodic parameters changed over most brain regions according to the task execution.

### No correlation between behavior or clinical features and aperiodic parameters

Contrary to our hypothesis we did not find any significant relationship between behavioral measures (RT, accuracy, congruence effect, first bin of CAF, last slope of delta plots) and aperiodic parameters. Literature provided mixed results about correlation between behavior and aperiodic activity. Some studies showed an association between the aperiodic parameters and cognition: Ostlund *et al*.^35^ found that a lower exponent during resting state predicted better performance on the dual-task “Stopping task”, Smith and colleagues^89^ showed that the aperiodic exponent was associated with cognitive performance and Ouyang *et al*.^90^ also evidenced that aperiodic activity during resting state was associated with cognitive processing speed. However, a study aiming to investigate aperiodic parameters and their link with cognition didn’t show any correlation between aperiodic activity and behavior.^86^ The absence of correlation between behavior and aperiodic parameters in our study was not expected based on the literature but there is currently a limited number of studies that have shown this association and mixed results highlight the importance of attempting to further investigate this potential link.

Despite significant changes in aperiodic parameters, those were not correlated to clinical characteristics of the PD population. This result is consistent with previous studies. Actually, no correlation between aperiodic parameters and UPDRS scores or motor-related sub-score have been found in the literature.^48,52^ Thus, this lack of clear association between aperiodic parameters and clinical criteria of the PD population suggests that aperiodic parameters may not be useful as clinical biomarkers. Another hypothesis is that the population of PD patients studied here did not experience sufficiently severe symptoms to reveal a clear association between clinical criteria and aperiodic parameters.

### Limitations

An important limitation of our study, which also applies to most PD-related studies, is that our PD population is different in terms of clinical criteria to PD populations in other studies, notably in age and disease severity, which can both influence cognitive control and EEG parameters. This limits our ability to compare our results and to establish firm conclusions. Regarding the methods, we used a brain template for source reconstruction, which limits the extent to which anatomical inter-individual variability is taken into account. We also used a specific brain atlas and it is not clear how the topography of aperiodic parameters may be impacted by using different atlas resolutions. Aperiodic components of the EEG have been linked to string subject-specific properties, which could explain at least in part the variability in our results.^91^ The small sample size used in our study also prevents the generalization of our results, motivating the validation on independent datasets.

### Perspectives

Several open questions remain with this study, notably the potential link between aperiodic parameters and cognitive alterations in PD. While our study did not provide support for an association between behavior and aperiodic activity, future efforts could include the use of other experimental cognitive tasks in which PD patients show alterations, such as a visual working memory task. Donoghue *et al*.^28^ have already shown that event-related changes in aperiodic parameters can predict individual working memory performance, therefore it could be tested whether or not this association holds in PD patients.

Furthermore, this study highlights the importance of taking into account aperiodic activity in EEG analyses, and provides further evidence that this component is linked to pathologies such as PD. In that sense, we suggest that studies focusing on the identification of clinical biomarkers of cognitive decline in PD using machine learning should consider aperiodic activity as an interesting feature.

## Acknowledgements

We thank all the participants of this study.

## Competing Interests

The authors report no competing interests.

### Abbreviations

BEM: Boundary Element Method
CAC: Cognitive Action Control
CAF: Conditional Accuracy Functions
DSM-V: Diagnostic and Statistical Manual of Mental Disorders - V
E:I: Excitation:Inhibition
EEG: Electroencephalography
FDR: False Discovery Rate
FIR: Finite Impulse Response
HC: Healthy Control
HD-EEG: High-Density EEG
ICA: Independent Component Analysis
ICBM MRI: International Consortium for Brain Mapping Magnetic Resonance Imaging
LEDD: Levodopa Equivalent Daily Dose
MoCA: Montreal Cognitive Assessment Scale
MSE: Mean Squared Error
PD: Parkinson’s Disease
PSD: Power Spectrum Density
REM: Rapid Eye Movement
ROI: Regions Of Interest
RT: Reaction Time
SNR: Signal to Noise Ratio
STN-DBS: Subthalamic Nucleus Deep Brain Stimulation
UPDRS: Unified Parkinson’s Disease Rating Scale
wMNE: weighted Minimum Norm Estimate

