## Supplementary Material for "Changes in electrophysiological aperiodic activity during cognitive control in Parkinson’s disease"

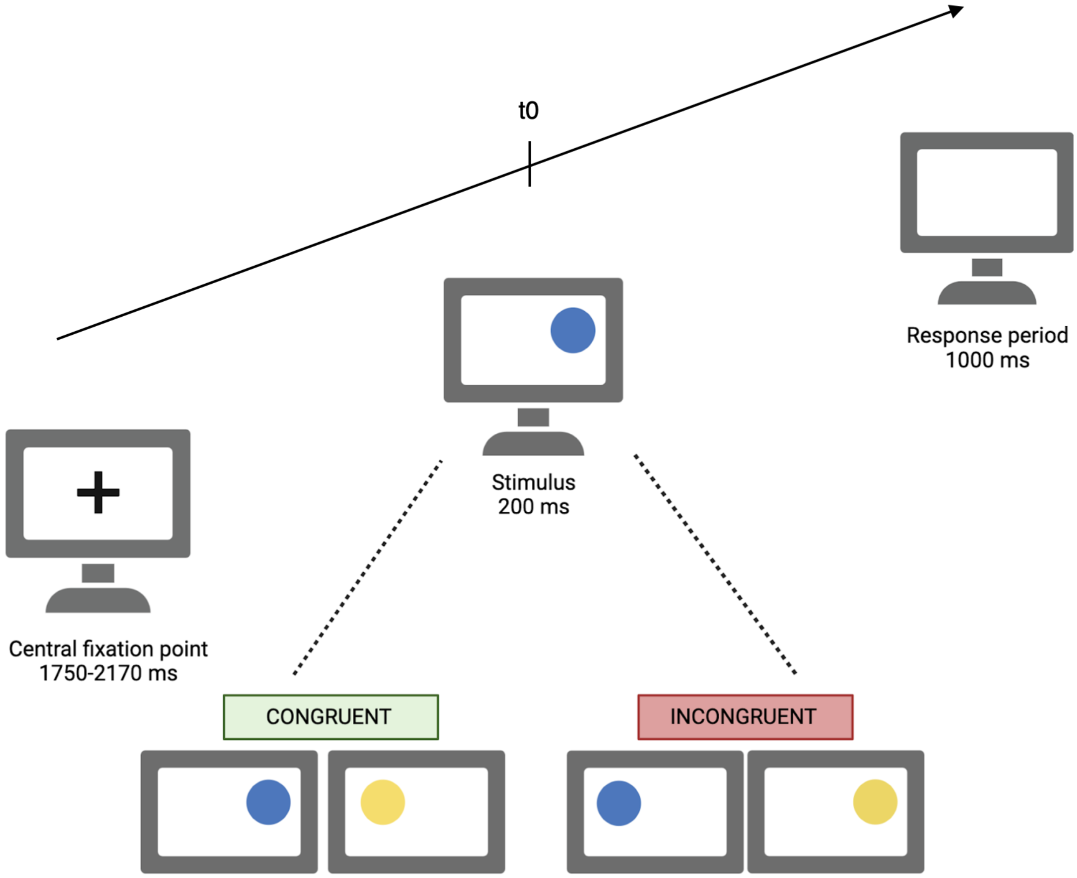


**Supplementary Figure 1 Overview of the Simon task design. Each trial began with a black fixation cross at the screen center during a randomly defined period between 1750 and 2170 ms.** Then, the stimulus was displayed during 200 ms. Participants had 1000 ms to answer by pressing the button. Two conditions could occur: congruent when the side of presentation of the circle and the required response side triggered the same response, and incongruent when both items did not lead to the same response.

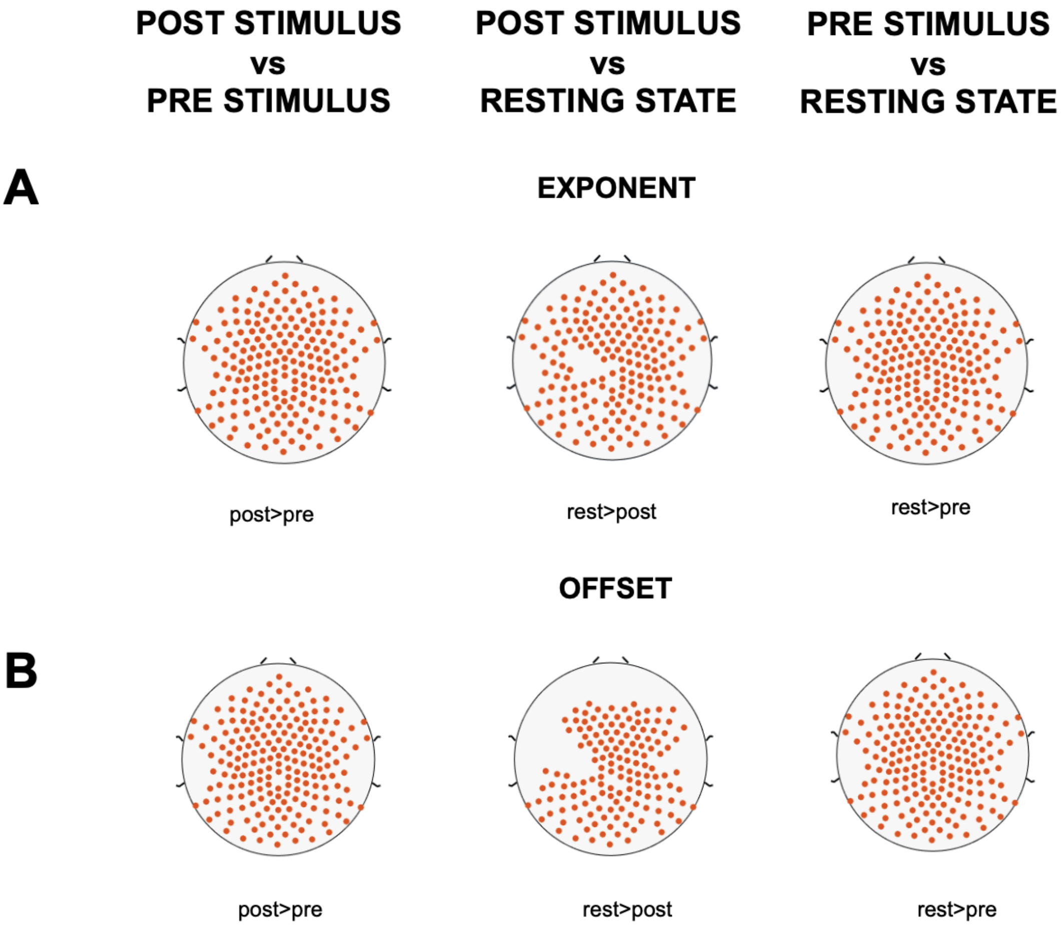


**Supplementary Figure 2 P-values distribution of t-tests (FDR-corrected) performed for each electrode across the scalp.** Comparisons of aperiodic exponent (A) and aperiodic offset (B) values were realized according to the task period. Electrodes with significant differences in aperiodic parameters between each period are represented with red dots.

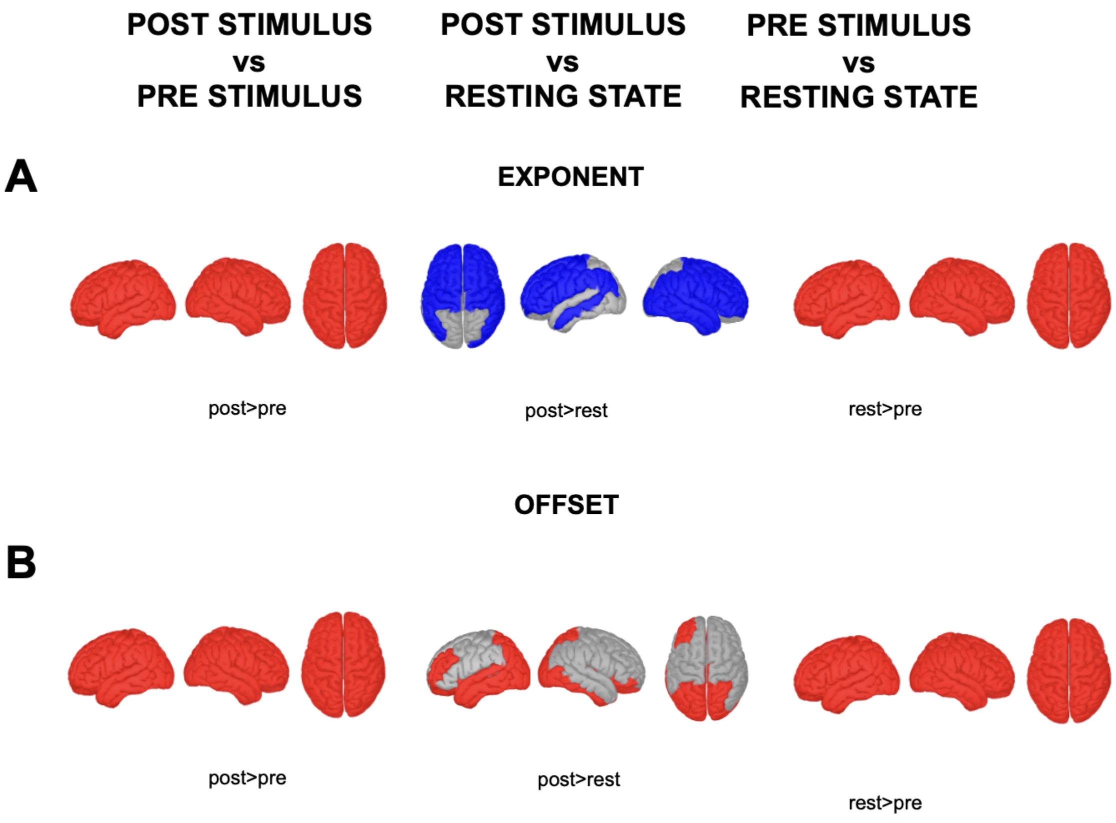


**Supplementary Figure 3 P-values distribution of t-tests (FDR-corrected) performed for each electrode across the cortex.** Comparisons of aperiodic exponent (A) and aperiodic offset (B) values were realized according to the task period. ROIs with significant differences in aperiodic parameters between each period are represented in color. Red color corresponds to significant differences in the same direction as the legend while blue color indicates significant differences in the opposite direction of comparison.

**Supplementary Table 1 Regions of interest where aperiodic parameters are significantly different according to the effect tested**

| Tested effect | Aperiodic parameter | Significant ROI |
|---|---|---|
| **Group – post stimulus part** | Exponent | - |
| | Offset | cuneus L and R<br>entorhinal L<br>fusiform L<br>fusiform R<br>inferior parietal L<br>inferior temporal L<br>lateral occipital L and R<br>lingual L<br>middle temporal L and R<br>parahippocampal L<br>pericalcarine L<br>postcentral L<br>precuneus L and R<br>superior parietal L<br>temporal pole L and R |
| **Congruence – post stimulus part** | Exponent | paracentral L and R<br>posterior cingulate L and R<br>precentral L and R |
| | Offset | paracentral L<br>posterior cingulate R |
| **Group – all parts** | Exponent | - |
| | Offset | bankssts L and R<br>cuneus L and R<br>entorhinal L<br>fusiform L and R<br>inferior parietal L and R<br>inferior temporal L and R<br>isthmus cingulate L and R<br>lateral occipital L and R<br>lateral orbitofrontal L<br>middle temporal L and R<br>paracentral L<br>parahippocampal L and R<br>pericalcarine L<br>postcentral L<br>posterior cingulate L and R<br>precuneus L and R<br>superior parietal L<br>superior temporal L and R<br>temporal pole L and R<br>transverse temporal L |
| **Task period** | Exponent | All |
| | Offset | All |

L: left; R: right.
